# Expanding the Potential Genes of Inborn Errors of Immunity through Protein Interactions

**DOI:** 10.1101/2021.05.10.443508

**Authors:** Humza A. Khan, Manish J. Butte

## Abstract

Inborn errors of immunity (IEI) are a group of genetic disorders that impair the immune system, with over 400 genes described so far, and hundreds more to be discovered. To facilitate the search for new genes, we need a way to prioritize among all the genes in the genome those most likely to play an important role in immunity. Here we identify a new list of genes by linking known IEI genes to new ones by using open-source databases of protein-protein interactions, post-translational modifications, and transcriptional regulation. We analyze this new set of 2,530 IEI-related genes for their tolerance of genetic variation and by their expression levels in various immune cell types.

## Introduction

IEIs are a collection of over 400 monogenic disorders with phenotypes of recurrent, severe or unusual infections, autoimmunity, and autoinflammation. The process of making a diagnosis in these patients requires the synthesis of the clinical phenotype, the results of immunological testing, and the results of genetic testing that identify the key pathogenic variant(s). Whole exome and genome sequencing have made the process of identifying genomic variants straightforward. But when this process cannot find a known pathogenic variant or even a known IEI gene, the process switches tracks to discovering whether a new gene might explain the disease phenotype. Paring down the list of thousands of genes and tens of thousands of variants in “non-clinical” genes/genes not yet identified as being important for a human disease, is an unsolved problem. The list of variants can be filtered to eliminate those variants that occur commonly in human populations, resulting in a shorter list of few hundreds to thousands of genes. However, beyond this step lies a painstaking process of choosing genes and testing their roles. Nowadays, over three dozen new IEI genes are discovered yearly [1]. Eventually, if the right gene has been identified, the ensuing process of validating the biochemical and immunological impacts of the identified variant(s) are well defined if arduous [2].

Here we describe a list of IEI-related genes that were gathered by associating known IEI proteins with new ones. A similar approach in 2015 created a list of over 3,000 IEI-related genes by linking 229 known (at the time) IEI genes to new ones by the human gene connectome (HGC) [3, 4]. The HGC was created by calculating genetic distance from the binding portion of the STRING protein database [5]. Expanding gene lists for other collections of rare diseases by such an approach has been fruitful [6]. The trade off with creating such a large list 3,000+ genes (more than 15% of the genome) is that many will be expected not to actually participate in non-redundant pathways of the immune response. Regardless, the list of known IEI genes since the 2015 paper has doubled, and the field needs an updated gene list.

In this work, we used the recently described OmniPath protein-interaction meta-database to reveal novel genes that are functionally related to IEI genes [7]. Specifically, we employed two routes to include putative IEI genes: 1) we analyzed annotated interactions encompassing transcriptional regulation, protein complex formation, and post-translational modifications (PTM) between two proteins, and 2) we analyzed pathways without functional annotation between all combinations of immunodeficiency genes present in the database.

## Materials and Methods

### Protein-Interaction Databases

We refer to Discovery pathway 1 as our method of collecting new genes by examining IEI genes with respect to transcriptional regulation, protein complex formation, and post-translational modifications (PTM) between two proteins. We refer to Discovery pathway 2 as our method of collecting new genes from all the pathways arising from all pairwise combinations of immunodeficiency genes present in the database.

OmniPath is a recently described meta-database of signaling pathways and protein interactions, integrating over 100 individual databases. We ran a list of 403 IEI genes into the OmniPathR package and infrastructure to derive the proteins most related to the relevant IEI gene products [8].

### Discovery Path 1

For analyzing transcriptional regulation, we imported interactions from the DoRothEA_A database [9]. IEI gene products were analyzed for their activity as transcription factors as well as their activity in inducing or repressing transcription of other genes. For post-translational modifications, we imported interactions from the SIGNOR database [10]. IEI gene products were analyzed for their activity as modifiers and as recipients of modifications.

### Discovery Path 2

Omnipath contains protein-protein association data from over 100 different databases. We queried whether a “pathway” could exist between all known IEI genes to other known IEI genes. If the Omnipath databases could identify a chain of protein interactions (a “pathway”) linking pairwise each known IEI gene, we then collected the genes along that pathway. To avoid the risk that every protein touches every other along some hypothetical pathway, we limited consideration to pathways with a total distance of less than 6; that is, an IEI gene would have to link to another IEI gene by less than six protein-protein interactions.

For IEI genes that did not return any interactions, we utilized the HGNChelper to find alternative gene symbols to query [11]. In sum, 334 IEI genes were analyzed for related proteins.

### Filtering pLI and GDI

We filtered pLIs under .9 using the updated pLI index available at https://gnomad.broadinstitute.org/downloads [12]. Gene damage indices (GDI) were taken from the HGC GDI server [13, 14].

### RNAseq Data

We used immune cell RNA sequencing data from the Human Blood Atlas [15]. This provided expression data from 18 different immune cell types (available at https://www.proteinatlas.org/humanproteome/blood). Monocyte subsets, eosinophils, basophils, and neutrophils were grouped as myeloid cells. Plasmacytoid and myeloid dendritic cells were grouped together as DCs. NK, B and T cell subtypes were grouped as such. Genes with an average of under 1 transcript per million in their group were filtered out of our group specific IEI candidate gene lists.

Genes present in our list of candidate genes that were not found in the RNAseq expression data (n=8) automatically were omitted from the filtered lists, and mostly included non-coding regulatory RNAs.

### Code

All plotting was done in ggplot2 in R. Code is available at github.com/humzalikhan.

## Results and Discussion

IEIs are genetic diseases, and every patient with a genetic disease deserves a specific genetic diagnosis whenever possible. The published IEI genes offer only a limited snapshot of all the human genes that underlie human disorders, as is shown by the rapid growth in the identification of new genes, more than 30 per year at this point [1]. We employed two methods in parallel to evaluate genes and include them into a list of potential IEI-related genes. First, we assessed protein-associations through the lenses of post-translational modification and transcriptional regulation. Second, we analyzed unannotated pathways of protein-protein interactions between all combinations of IEI genes (Fig. 1). Together, these approaches identified a list of 2,530 genes that we propose should be prioritized for consideration as IEI genes (Table S1). The known IEI genes have been split into nine categories of immune function (“tables” in the IUIS paper [1]) (Fig 2A). Our list of 2,530 genes associated more with Table 1 genes that cause cellular and humoral defects, and almost none with genes that cause complement deficiencies (Fig. 2B).

**Figure 1.**
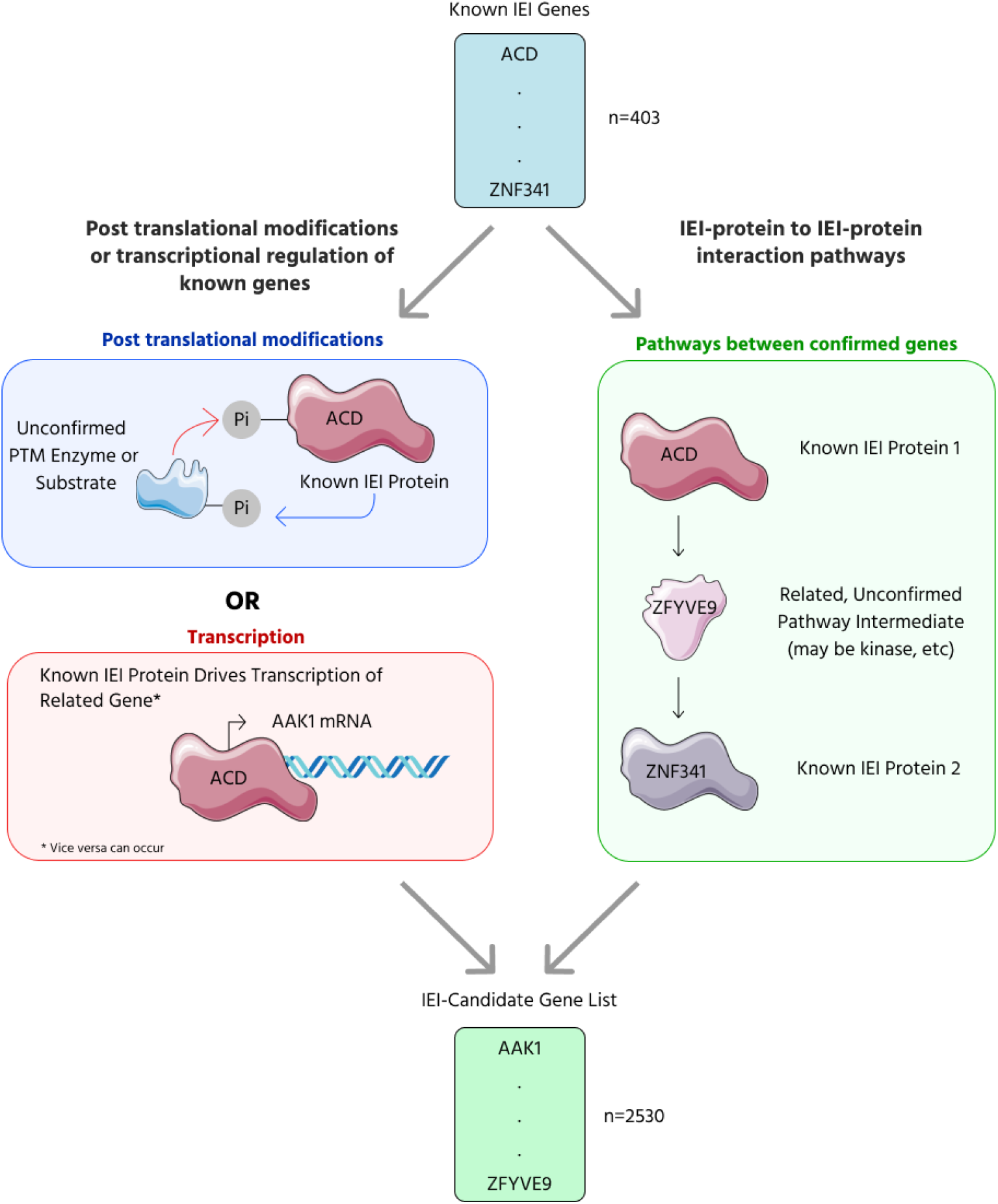
Overview of our approach. We used the OmniPath set of databases to create a list of candidate IEI genes using two approaches: pathway 1 (left), which uses post-translational modification and transcription factor data, and pathway 2 (right), which uses protein-protein interaction pathways.

**Figure 2.**
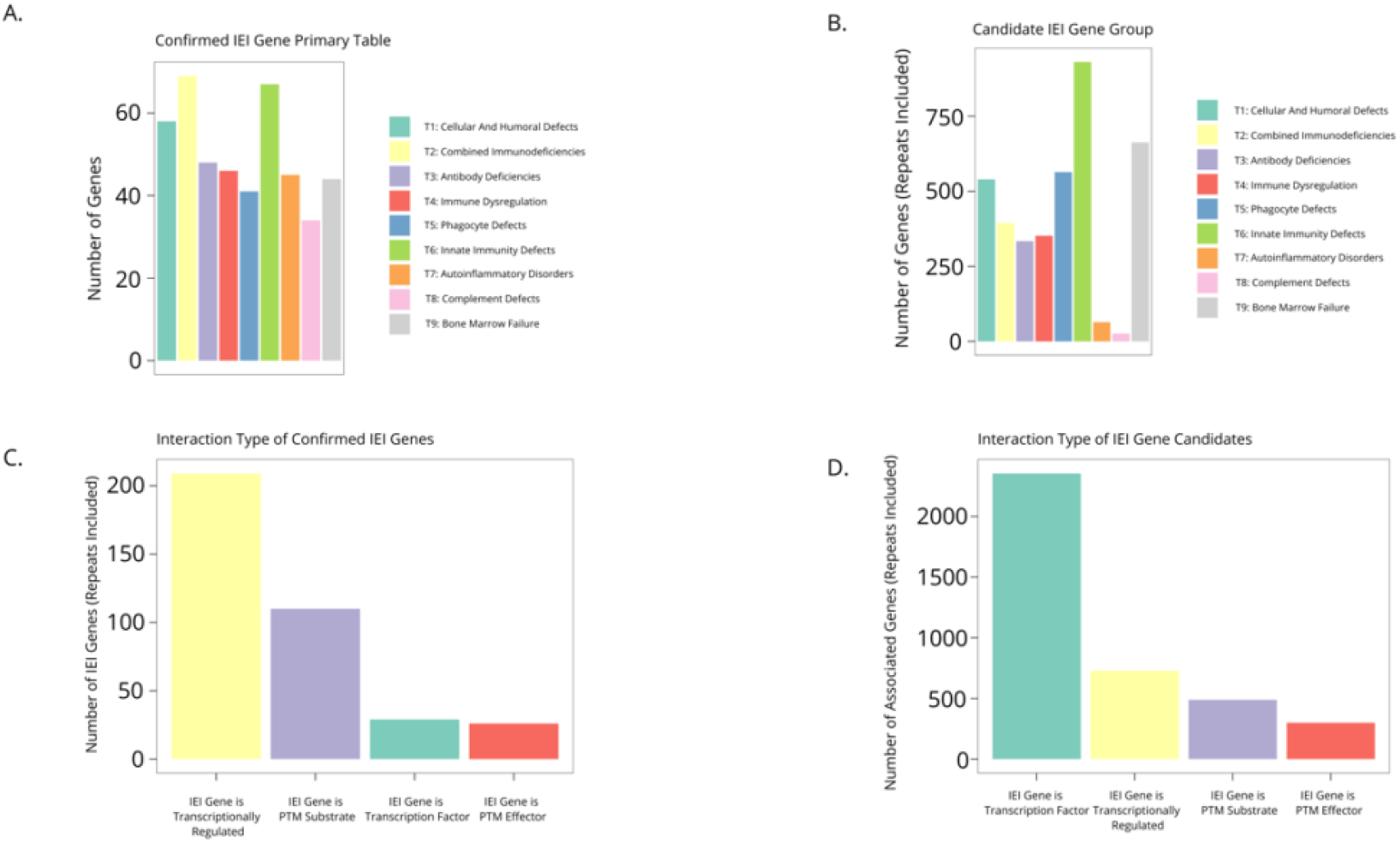
Discovery Pathway 1 reveals IEI candidates by analyzing transcriptional regulation and post-translational modifications of known IEI genes. **(A)** Number of confirmed IEI genes in respective IUIS tables as defined by the phenotype that their mutations cause. **(B)** Number of candidate IEI genes in tables classified by their interactions with known IEI genes. For A and B, only primary tables were utilized. **(C)** Number of known IEI genes and their respective modifications. **(D)** Number of candidate IEI genes performing transcriptional regulation on IEI genes, being regulated by IEI genes, and PTMs to/from IEI genes.

In our pathway 1 approach, we analyzed 334 IEI-related genes out of the 403 of the IUIS IEI gene list. Each gene was analyzed for its action as a transcription factor, its induction/repression due to a transcription factor, and activity as a post-translational modifier or as a recipient post-translational modification. Of the relationships described in pathway 1, we found that IEI gene products are post-translationally modified more often than they post-translationally modify other proteins (Fig. 2C, 2D). In particular, they are likely to be phosphorylated along a signaling pathway which results in cellular response such as transcription. Some IEI genes also encode kinases/phosphatases themselves (Fig. S1A). While IEI gene products also ubiquitinate and cleave other proteins, it was found that modifications to IEI genes are more diverse than the effects of known IEI genes. Associated genes may (de)methylate, neddylate, de(acetylate), and sumoylate IEI genes. Some of these interactions have been described in literature; for example, SUMO modification of STAT1 has been well-documented [16], but the sumoylation enzymes are not known IEI genes. As the discovery of the *TNFAIP3, OTULIN, HOIL-1*, and *HOIP* have shown, PTMs are important to regulate proper immunity activity and quiescence; conceivably, a deleterious variant in the genes that modify or are modified by known IEI proteins could alter immune function and cause a novel immunodeficiency.

In discovery pathway 1, we also analyzed transcriptional regulation related to IEI genes. Notably, we found that IEI proteins more often are involved as effectors of this regulation as opposed to recipients of it. The STAT family is a set of transcription factors of many downstream pathways relevant to immune function; defects in STAT action are known to cause susceptibility to infection, autoimmunity and immune dysregulation [17]. Defects in the downstream products of IEI genes that act as transcription factors (TFs) and the upstream TFs that induce transcription of IEI genes may alter immune function and be found to cause immunodeficiency.

In pathway 2, we constructed a set of putative “pathways,” the starts and ends of which are every known IEI-gene. Along the pathway lie validated protein-protein interactions, validated by literature. For example, two proteins that lie along the pathways between IFNAR1 and STAT3 include TYK2 or JAK1. Alternatively, longer, less biologically probable routes exist, such as WRAP53 -> TP53 -> RPS6KA4 -> MAPK14 -> CSNK2A2 -> DKC1. In discovery pathway 2, we limited the gene-gene pathway distance to five to reduce the biological irrelevance of the putative pathways, knowing that six or seven degrees of separation link virtually every protein with every other. Our analyses of protein-protein interaction pathways from known IEI gene to known IEI gene revealed many previously known IEI genes (Fig 3A), a result supports that as-yet undiscovered IEI genes may lie along these same routes. We found many new genes using this approach (Fig 3B). We again found complement-related genes underrepresented, similar to the functional annotations in discovery pathway 1. This result suggests that not many pathways exist ending in a complement protein, that the complement system is perhaps independently regulated and is less intertwined with other immune defenses [18].

**Figure 3.**
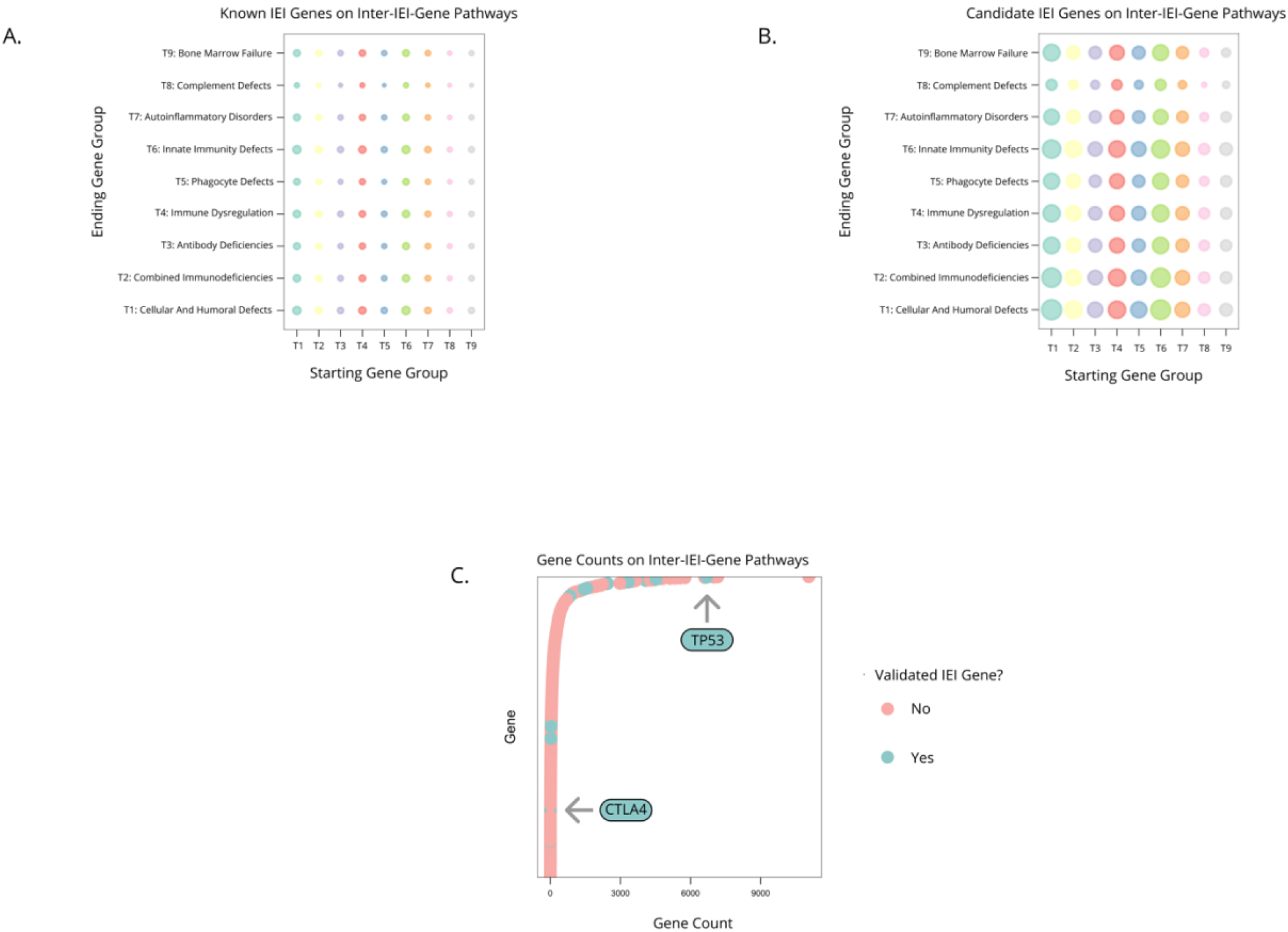
Discovery Pathway 2 uses protein interaction data to describe pathways between genes. **(A)** Bubble plot showing the relative amount of known IEI genes lying on IEI gene to IEI gene protein interaction pathways. **(B)** Bubble plot showing the relative amount of candidate IEI genes lying on IEI gene to IEI gene protein interaction pathways. **(C)** Count of genes on IEI gene to IEI gene pathways, with previously known IEI-causative genes indicated.

To assess whether a particular gene might play a role in multiple pathways, and thus potentially be important to multiple mechanisms of disease, we analyzed the frequency of genes along the protein-protein association pathways. However, many known IEI genes occurred at high frequency within these pathways, while others only occurred one or a few times (Fig. 3C). For example, CTLA-4, a well-known inhibitory receptor whose deficiency causes antibody deficiency and autoimmunity, was only found once as a pathway intermediate in between two IEI genes. Therefore, we decided against culling low-frequency genes from our list of candidates.

Our list of 2,530 related genes cuts down from the previously described list of over 3,000 candidates from the Itan group [3]. Furthermore, since we utilized a compendium of many databases, our associations are likely to be biologically relevant and supported by literature. About half of the genes we identified appeared in the Itan group’s list, and the other half were novel.

Ranking genes by their probability of loss intolerance (pLI) has been suggested as a way to predict the importance of unknown genes to human disease, especially for genetically dominant conditions. High pLI values imply purifying selection in the population when genomic loss-of-function variants (e.g., frameshift, truncation) appear. pLI values are calculated empirically from large databases of healthy individuals by comparing observed variants in the population to the number of variants expected to arise by chance [19]. These databases and pLI values are of course subject to change as more and diverse populations are sampled. To filter our putative list of IEI genes, we considered pLI scores for all known and putative IEI genes. Notably, we found that known IEI genes did not have unilaterally high pLI (Fig. S2A). Disaggregating by inheritance type, high pLI values (>0.9) were shown to be associated with autosomal dominant inheritance and well associated with X-linked dominant conditions. The pLI for known IEI genes is hardly constrained for autosomal recessive disorders (Fig. S2A). Thus, we have provided a shorter list of 810 high-pLI genes (including genes without available pLIs) for consideration of X-linked dominant or autosomal dominant conditions (Table S2). This subset of genes may be less important for recessive clinical traits, where the entire list of putative genes should be considered.

Since not all IEI genes have high pLI scores, we decided to explore the previously described Gene Damage Index (GDI) meant to predict the deleterious capacity of a variant in a gene [12]. However, we found that many known IEI genes also did not score highly on the predicted GDI (Fig. S2B). Predictions of damage from known IEI genes using the GDI results in mostly a “medium” characterization (Fig. S2C). Therefore, qualitative and quantitative GDI predictions have limited utility for filtering putative IEI genes. We found that GDI and pLI do not correlate with each other in known IEI genes (Fig. S2D). Taken together, our results show that filtering on only high pLI or high GDI genes would serve only a limited role and may unnecessarily constrain lists of putative genes.

Known IEI genes are generally expressed in immune cells; we used RNA sequencing data from the Human Protein Atlas to liberally filter for variants expressed in any immune cell type. The Human Protein Atlas database includes RNA sequencing data of 18 immune cell types from healthy donors which comprised five general groups: B cells, dendritic cells, myeloid cells (basophils, neutrophils, eosinophils, monocytes), NK cells, and T cells [15]. For initial inquiry, we analyzed the expression of all genes in each cell group and found most genes not expressed at all in immune cell types (Fig. S3A). However, when only IEI genes were plotted, the distribution was more right-skewed, indicating the expected trend that IEI genes are transcriptionally well expressed in immune cells (Fig. S3B). Expectedly, among the IEI genes that have zero RNA transcripts among immune cells are AIRE and complement genes. Therefore, we decided to filter a list for expression above 0 TPM in any immune cell type (Table S3).

Furthermore, since many IEIs manifest with cell-type specific defects, we decided to use RNA sequencing data to also create IEI-related gene lists based on cell-type specific gene expression. Such a list may offer utility when investigators are focused on disordered with focused immunophenotypes. For example, one might begin a search for new SCID genes where only T cells are affected by looking at transcripts that are expressed in T cells or T cell precursors.

To determine a cutoff for what is considered “expressed,” we plotted gene expression of IUIS table 1 genes, which affect humoral and cellular immunity, and table 2 genes, which affect just antibody production, and disaggregated by cell type. The former would be more likely to be expressed in all lymphocytes and the latter in only B cells (Figs. 4A, B). As disorders in table 1 can manifest due to defects in either T or B cells, we found that many of the genes that were expressed at a low level in T cells were expressed in B cells and vice versa. For example, among the low-expressed genes in B cells, for example, was *CD3D. CD3D* is a major component of the T-cell receptor complex and was highly expressed in T cells but not in B cells. CD3D deficiency causes a T-/B+ SCID (a table 1 condition) and is thought to be irrelevant in B cell development, which is supported by its negligible gene expression. Thus, if one were considering a list of IEI genes to be used for patients with B-cell focused defects, we would filter out such T-cell-specific genes as likely irrelevant.

**Figure 4.**
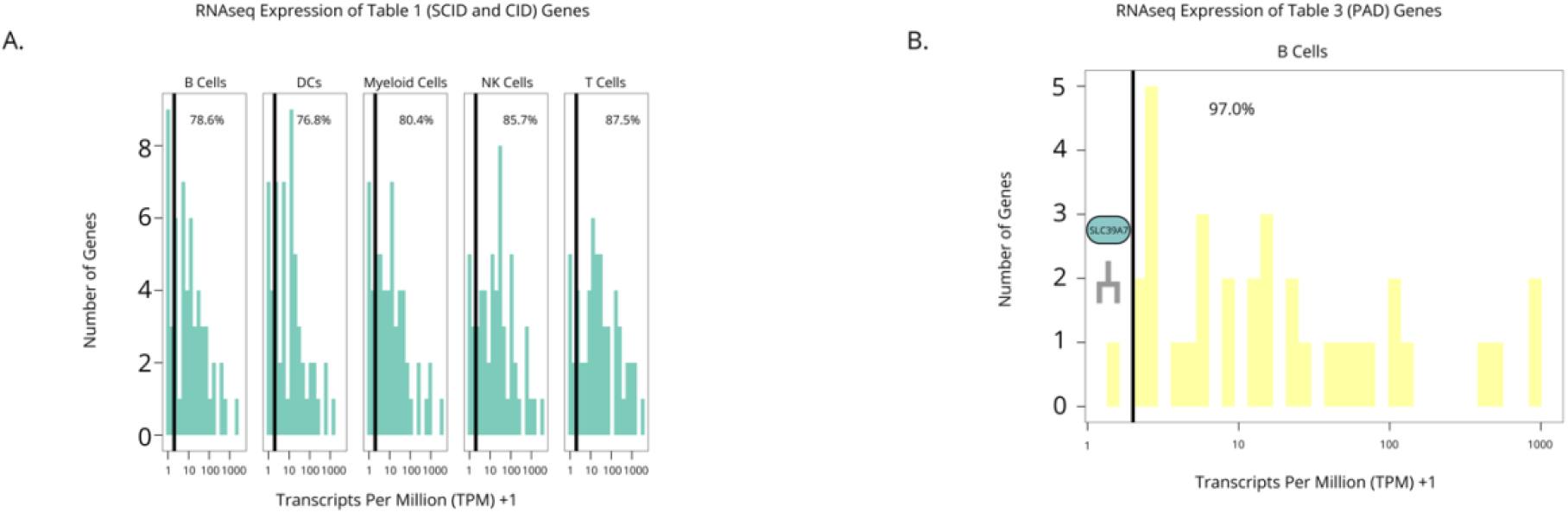
RNA sequencing data provides a method to filter for relevant IEI genes dependent on patient phenotype. **(A)** Expression levels of SCID and CID-causing genes in the cell types found in the Human Blood Atlas RNAseq expression matrix. **(B)** Expression levels of antibody deficiency-related genes in B cells. Transformations of TPM+1 done to visualize 0 values. Percentage above 1 TPM (at 2 when transformed to TPM+1) cutoff presented in each plot.

We set our cutoff point for expression as greater than 1 transcript per million (TPM). We picked this limit based on the transcriptional expression of known IEI genes across cell types (Figs. 4A, B). There was only one known IEI gene that causes a primary antibody deficiency (PAD) expressed below 1 TPM in B cells, the gene called *SLC39A7*, which codes for ZIP7. Ostensibly, this finding was unexpected since PADs reflect defects in B cell numbers or function, but an obvious explanation would be that this gene is expressed only early in B-cell development and not in peripheral B cells (Fig. 4B). Indeed, ZIP7 is an essential zinc transporter without which B-cell development is abrogated in the transition from pro- to pre-B cell. Thus, we anticipate that filtering gene lists by expression levels would preserve the majority of interesting genes. The genes that were erroneously filtered out in cell-type specific lists would still be present in the broader, unfiltered list. Our cell-type specific lists for the cell type groups in the Human Blood Atlas are found in the supplement (Tables S4-8).

A limitation of our work is that some genes are not described in the protein-interaction databases queried. For example, the complement protein C2 with its known interaction with CD19 was not represented in the protein-protein databases, and this interaction was thus not recognized in our searches using the OmniPath databases. Furthermore, the Human Blood Atlas database holds RNAseq data on major categories of immune cells; gene expression data from clinically relevant subsets such as plasmablasts, effector memory T cells, and others would be valuable. As access to open-source datasets increases and as functional evidence for protein function is released, we will refresh our lists.

In summary, by using a combination of both 1:1 annotated protein interactions and larger, un-annotated protein-interaction pathways, our approach allows for both a global and local view of proteins that may be relevant to query in future immunodeficiency studies. Notably, we found that high pLI or GDI were not particularly good criteria for determining the pathogenicity of a putatively novel IEI gene, especially those with recessive patterns of inheritance. Our work further advances on previous studies by merging transcriptional expression data with our list of IEI candidate genes derived from protein interactions to ensure that queries based on clinical presentations (i.e., T-cell lymphopenia) or diagnostic hypotheses can be made.

## Author Contributions

Conceptualization, Analysis, Writing: H.A.K. and M.J.B.; Funding Acquisition, Project administration, Supervision: M.J.B.

## Data Availability

All data for this study are included in the manuscript and the supplementary files.

