## Supplementary Figures for "Expanding the Potential Genes of Inborn Errors of Immunity through Protein Interactions"

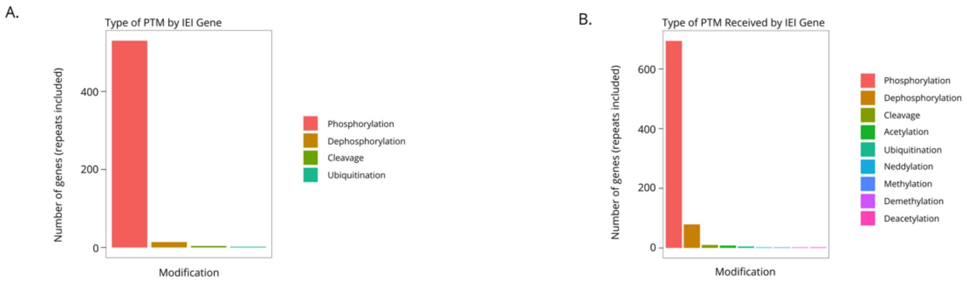


**Figure S1. Post-translational modifications induced by and received by IEI gene products.**

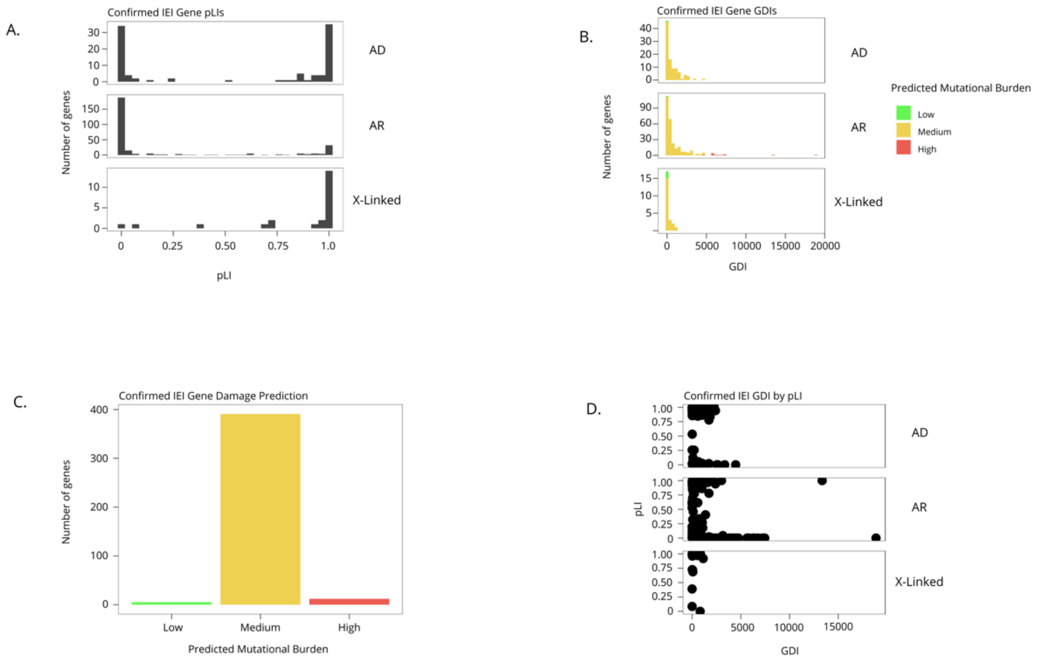


**Figure S2. Known IEI-causative genes have varying levels of predicted mutational harm.** **(A)** Confirmed IEI gene pLI disaggregated by inheritance type (AD: autosomal dominant, AR: autosomal recessive). **(B)** Confirmed IEI gene Gene Damage Indices (GDIs) disaggregated by inheritance type. **(C)** Damage prediction of confirmed IEI genes as classified by the GDI Server. **(D)** Known IEI gene pLIs plotted against GDIs and disaggregated by inheritance type.

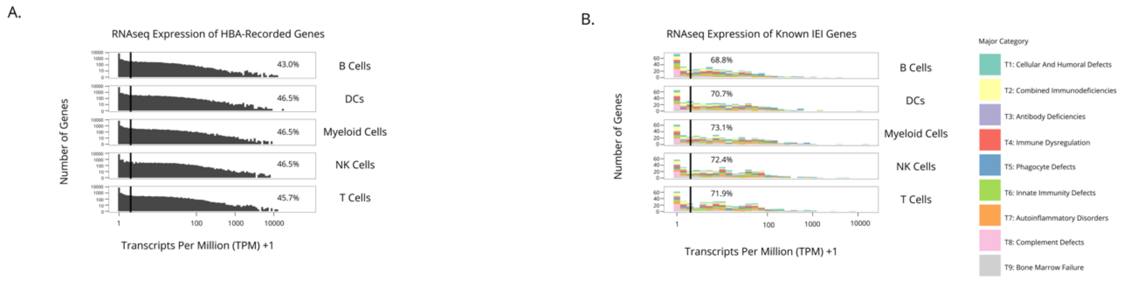


**Figure S3. Known IEI transcripts skew to higher expression in immune cell types. (A)** RNAseq expression of all Human Blood Atlas-recorded genes disaggregated by cell types present. **(B)** RNAseq expression of all IEI genes disaggregated by cell type. Line drawn at 1 TPM (at 2 when transformed to TPM+1). Percentage above 1 TPM (at 2 when transformed to TPM+1) cutoff presented.
